# A formal model of anxiety disorders based on the neural circuit dynamics of the fear and extinction circuits

**DOI:** 10.1101/2025.01.20.633839

**Authors:** Alan Lawrence Rubin, Mark Walth

## Abstract

The pathophysiology of anxiety disorders is the outcome of an imbalance of the fear-anxiety circuit and the extinction circuit. We present a formal model using nonlinear dynamics and network theory, which captures the dynamic interactions of the key nodes of the anxiety and extinction networks. This rudimentary model can be modified by newer data. These core nodes consist of the cells of the paraventricular nucleus of the thalamus coding negative valence, the neurons of the basal-lateral amygdala coding negative valence (Rspo2+), the anterior cingulate cortex, the ventral hippocampus neurons coding fear memories, the somatostatin expressing cells of the lateral segment of the central amygdala, the medial segment of the central nucleus of the amygdala-bed nucleus of the stria terminalis and their target nodes. The extinction network consists primarily of the paraventricular thalamic cells coding positive valence, (Ppp1r1b+) cells of the basolateral amygdala coding positive valence, the ventromedial prefrontal cortex, the PKCδ cells of the lateral segment of the central amygdala, and the intercalated cells.

Human and non-human animal genetic and epigenetic studies point to deficiencies in brain-derived neurotrophic factor and neurotrophic receptor kinase tyrosine 2 production in key nodes causing reduced plasticity extinction network plasticity and leading to a weakened extinction response.

We rely primarily on the neurophysiological studies of non-human animal models since nodes generating fear/anxiety and extinction responses are highly conserved across species and equivalent nodes are present within analogous circuits of the human brain. The results are confirmed, where possible by human functional magnetic resonance imaging studies.

We believe this simplified model is of heurist value and can lead to a more consistent focus on physiologically based pathophysiology. This would lead to treatments to reverse the pathologic physiology produced by genetic and epigenetic abnormalities, and greater efforts to directly correct pathologic circuit activity through direct interventions such as transcranial magnetic stimulation.

**Significance:** We believe this simplified model is of heurist value and can lead to a more consistent focus on physiologically based pathophysiology. This would lead to treatments to reverse the pathologic physiology produced by genetic and epigenetic abnormalities, and greater efforts to directly correct pathologic circuit activity through direct interventions such as transcranial magnetic stimulation.

## Introduction

This paper we assume that fear and anxiety have the same neurophysiological underpinning and are, therefore, physiologically identical. Brain centers producing fear and its extinction form opposing circuits-the fear and fear extinction circuits [1–4]. Using network theory, we can understand the pathophysiology of anxiety disorders as the outcome of an imbalance between the strength of output of the fear-anxiety circuit-reacting to a stimulus assigned a negative valence- and an extinction circuit reacting to a competing positive valence stimulus [5–12].

Our model of neural interactions is primarily based upon animal studies, since the nodes for generating fear/anxiety responses are highly conserved [14–22] Specifically, the rat infralimbic cortex (IL) is believed to be a rudimentary form of the human ventromedial prefrontal cortex (vmPFC), (Brodmann”s area 25) and the rat prelimbic cortex is considered an equivalent of the human anterior cingulate cortex (ACC) (Brodmmann area 32) due to similarities in connectivity and role in fear consolidation [18–22]. When possible, we obtain supplementary evidence from human fMRI studies though fMRI does not currently have the degree of resolution necessary to precisely identify nodal activity [23].

### Description of the nodes comprising the fear and extinction circuits

The fear circuit includes nodes that originate and amplify the fear signal and those that execute physiological and behavioral fear responses. The nodes producing an initial negative valence signal are cells of the paraventricular thalamic nucleus (PVTfear) and basolateral amygdala R-spondin 2-expressing (Rspo2+) neurons (BLAfear). The BLA negative valence signal is passed to the lateral somatostatin expressing cells of the central amygdala – the CeLon

Cells of the ventral hippocampus (vHPC) coding negative valence memory engrams (vHPCfear) have recurrent connections to BLAfear. Fear recall is associated with increased c-Fos expression-a measure of heightened neural activity-in vHPC projections to the BLA in humans. The weighting of the fear-anxiety signal is further heightened by recurrent connections of the anterior cingulate cortex (ACC) with BLAfear neurons [5–11,29–32,39]. The extinction network consists primarily of the PVT cells coding positive valence (Ppp1r1b+), cells of the BLA coding positive valence, the ventromedial prefrontal cortex (vMPFC), the PKCδ cells of CeL, and the intercalated cells (ITC). BLA (ext) receives input from vHPC (ext) coding extinction memories. BLA (ext) stimulates CeL PKCδ cells (CELoff) which inhibit CeM-BNST and thereby dampen sympathetic arousal. IL and BLAext activate the intercalated cells (ITC) which also inhibit CeM-BNST [5–12,24–38,40].

The vmPFC is a key extinction center. Extinction recall is associated with increased activity of vHPC projections to the infralimbic cortex (IL). In humans, the level of vmPFC activity is directly proportional to the degree of extinction recall. vmPFC inhibits BLAfear: Control subjects show greater activity of the vmPC than subjects with anxiety disorders during fear inhibition and have greater vmPFC connectivity to the amygdala (AMYG). Increased activity in the vmPFC is associated with more positive valence ratings. Human high rest anxiety is negatively correlated with amygdala– vmPFC functional connectivity on fMRI [41–46].

Both fear/anxiety and extinction begin with the assignment of a negative or positive valence through the input of PVT to BLA. It is possible that the valence of the PVT neurotensin afferents determines the valence of each targeted BLA neuron [24-27

### Genetic and epigenetic changes in extinction network nodes cause anxiety disorders

BDNF and its receptor neurotrophin receptor kinase 2 (NTRK2) are necessary for the growth of synaptic connections leading to synaptic plasticity and increased synaptic communication [61–63]. Genetic variants such as the BDNF allele ValMet and stress-induced methylation of the genes for BDNF and its receptors lead to deficiencies in BDNF function [64–71]. BDNF deficiencies caused by genetic variants or epigenetic changes occurring in the key nodes of the extinction system-the vmPFC, AMYG and VHPC can increase anxiety by impairing theextinction response [72–79, 83–87]. The accumulation of fear reactions without sufficient counterbalance by a functioning extinction network leads to anxiety disorders [80–82].

## Method

We construct a model based upon the likely activity of the key nodes of the fear and extinction networks. Using network theory, we can conceptualize the vast complexity of the brain as an array of nodes connected by large bundles of myelinated tracts or edges. Nodes with complementary functions connect within larger networks in a hierarchical manner with each later node refining the output of the prior [88,89]. Mathematical formulations can offer a precise foundation for understanding the interactions of brain centers and their interconnections causing anxiety disorders.

We must first identify and justify the underlying assumptions of the formal model for anxiety disorders. These assumptions are:

1. Anxiety disorders are disorders of neural circuits.
2. Based on current research the most likely circuits are the fear/anxiety circuit and the extinction circuit.
3. The activity of each network is dependent on the activity of each of its components.
4. Anxiety states are not attributable to a specific node but arise from the combined interaction of individual nodes tied to their circuits and the interactions of those circuits.
5. The symptoms of anxiety disorders arise from dysfunction within networks and between interacting networks caused by an imbalance in the plasticity of new synapse production between the fear/anxiety and the extinction networks.
6. The interactions of the networks are best quantified by using the nonlinear dynamics of neuron firing rates.
7. The firing rate of each node obeys the so-called *firing rate differential* equation, a system of coupled ordinary differential equations. [13,88]

In our model, the fear generating system ascribes negative valence to specific sensory experiences or memories and produces the array of psychological, behavioral and physical symptoms associated with fear/anxiety.

### Network Model

The dynamics of the system are modeled by coupled ordinary differential equations describing the evolution of the firing rate of each node. There is one equation for each node of the graph. The interconnections of the network are modeled as a weighted directed graph, with weightings representing the strength of the connections within and between the individual nodes in the network. These weightings appear in the governing differential equations in the form of the weighted matrix *W*.

Network activity arises from a relative firing rate. The strength of the signal communicated between nodes of a circuit or by the entire circuit is dependent on firing rate, synaptic density and sensitivity which is assigned a weighting (*W*). This activity would be averaged over periods of hours, days, weeks or months, typical of the variety of time lengths for episodes of the different anxiety disorders.

Positive and negative weightings indicate excitatory or inhibitory connections. We use the convention that *W*ij denotes the influence that node i has on node j. For example, if PVTfear is assigned #1 and BLAfear is assigned #2, then *W*1,2 denotes the influence of PVTfear on BLAfear.

We use weightings to produce levels of network activation that model the physiological variables primarily produced by animal models and when possible, fMRI results of network nodes associated with the specified anxiety disorders. In our model all network variables must be relative since the actual values for average frequency, weightings, and slope of activation in anxiety states are not known.

Circuits modeling each anxiety disorder are assigned a relative weighting as a fraction or multiple of its weighting in the resting state. We assign weightings to each input of a circuit based upon neurophysiologic models of the activity of each node in nonhuman animal models of fear and extinction. In anxiety states, nodes of the fear circuit are assigned higher values than in the resting (non-anxious) state and nodes of the extinction circuit are assigned lower values

### Formulation of network equations

In the formal model, the strength of influence of unit 𝑖 on unit 𝑗 wi is represented by a scalar called the *weighting,* which will be denoted 𝑤_𝑖𝑗_. The governing equation of such a firing rate network is given by equation 7.10 of [13]:

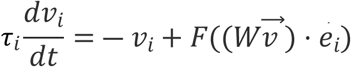

where:

- 𝑣_𝑖_ is the firing rate of unit 𝑖. For example, 𝑣_𝑆_ denotes the firing rate of the salience network.
- 𝑣 is a vector containing the firing rate of each unit, 𝑣 = (𝑣_𝑆_,𝑣_𝐸_,𝑣_𝐷_);
- 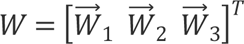 is the matrix containing the weightings of all the connections within the network, and 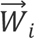 denotes the weightings of all connections going into node 𝑖;
- 𝜏_𝑖_ is a time constant.
- 𝑒_𝑖_ is the 𝑖^𝑡ℎ^ standard basis vector.
- 𝐹 is the “activation function” of the neurons, representing how the firing rate of a neural unit changes as a function of its input current.

We use the sigmoidal activation function:

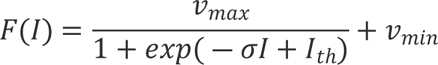 In this expression, 𝐼 is the total input current to the neural unit, 𝑣_𝑚𝑎𝑥_ is the maximum firing rate of the neural unit, 𝑣_𝑚𝑖𝑛_the minimum possible firing rate - which we take to be small but nonzero - 𝐼_𝑡ℎ_ is the activation threshold, and 𝜎 is a parameter which controls the steepness of the activation curve. Biologically, the parameter 𝜎 represents the overall sensitivity of a neural unit to input current. In a biological neuron, sensitivity is determined by a range of factors, including ion channel density and synaptic density and sensitivity. In some texts, see for example [13], the constants are taken to be 𝑣_𝑚𝑎𝑥_ = 100Hz, 𝐼_𝑡ℎ_ = 50nA, 𝜎 = 17.5, and 𝜏 = 5ms. We choose to scale our parameters in such a way that emphasizes qualitative behavior of the network, rather than focusing on specific quantitative relationships.

In this scaling, 𝑣_𝑖_represents the firing rate of unit 𝑖, represented as a fraction of its maximum possible firing rate. For example, if 𝑣_𝑖_ = 0.6, then unit i is firing at 60% of its maximum possible firing rate. Finally, for simplicity, we take all time constants to be equal and set them to 𝜏_1_ = 𝜏_2_ = 𝜏_3_ = 5. Therefore, the parameter that we manipulate in this model are the weightings between neurons, 𝑊.

In this paper we assume anxiety disorders result from deficiencies of BDNF and its receptors in key nodes of the anxiety extinction network. Our equations of network functio model a paucity of BDNF, which causes a decrease in synaptic weighting due to a decrease in synaptic density, decreased amplitude of excitatory post synaptic current (epsc) and reduced quanta of transmitter release.

## Results

We apply a formal dynamic neural circuit model to our construct of the fear and extinction circuits derived primarily from nonhuman studies. The goal is to demonstrate how alterations in the weightings of connections between key nodes of the fear and extinction circuits may produce common anxiety syndromes. We favor the data produced by physiological studies in non-human subjects because the data is accurate, finely detailed and sufficiently replicated to construct a working neurocircuit model. Where possible we include supporting data from human fMRI studies.

We assume that nonhuman findings pertain to human anxiety disorders especially for those disorders that share bot similar circuits and similar symptoms: generalized anxiety disorder, panic disorder and post-traumatic stress disorder.

For the resting state all within-circuit weightings between nodes of the fear-anxiety circuit and the extinction circuit are equivalent. All weightings are set at (+1) for excitatory input and at (−1) for inhibitory input at the baseline state. The resulting stimulation of the predominant output nodes of the fear circuit-BLAfear, vACC, vHPC and CM-BNST are at a low level. Lowering BLAfear causes low stimulation of vHPC fear, and ACC. Low CM-BNST activation causes low stimulation of PAG, hypothalamus, and LC.

Nodes of fear circuit are: 1 =PVTfear, 2 = BLAfear, # 3 = vHPCfear, # 4 = ACC, #5=CeLon. Nodes of extinction are: # 6 =PVText, # 7 =BLAext, #8=vMPFC, # 9 = vHPCext, # 10 - CeLoff, # 11 - ITC, # 12 = CeM/BNST.

Weightings for the resting state are: W[1,2] = 1, W[1,5] = 1, W[2,3] = 1, W[2,4] = 1, W[2,5] = 1, W[3,2] = 1, W[4,2] = 1, W[5,10] = −1, W[5,12] = 1, W[6,7] = 1, W[7,10] = 1, W[7,11] = 1, W[8,2] = 1, W[9,8] = 1, W[10,5] = −1, W[10,12] = −1, W(11,12] = −1

**Fig. 1.**
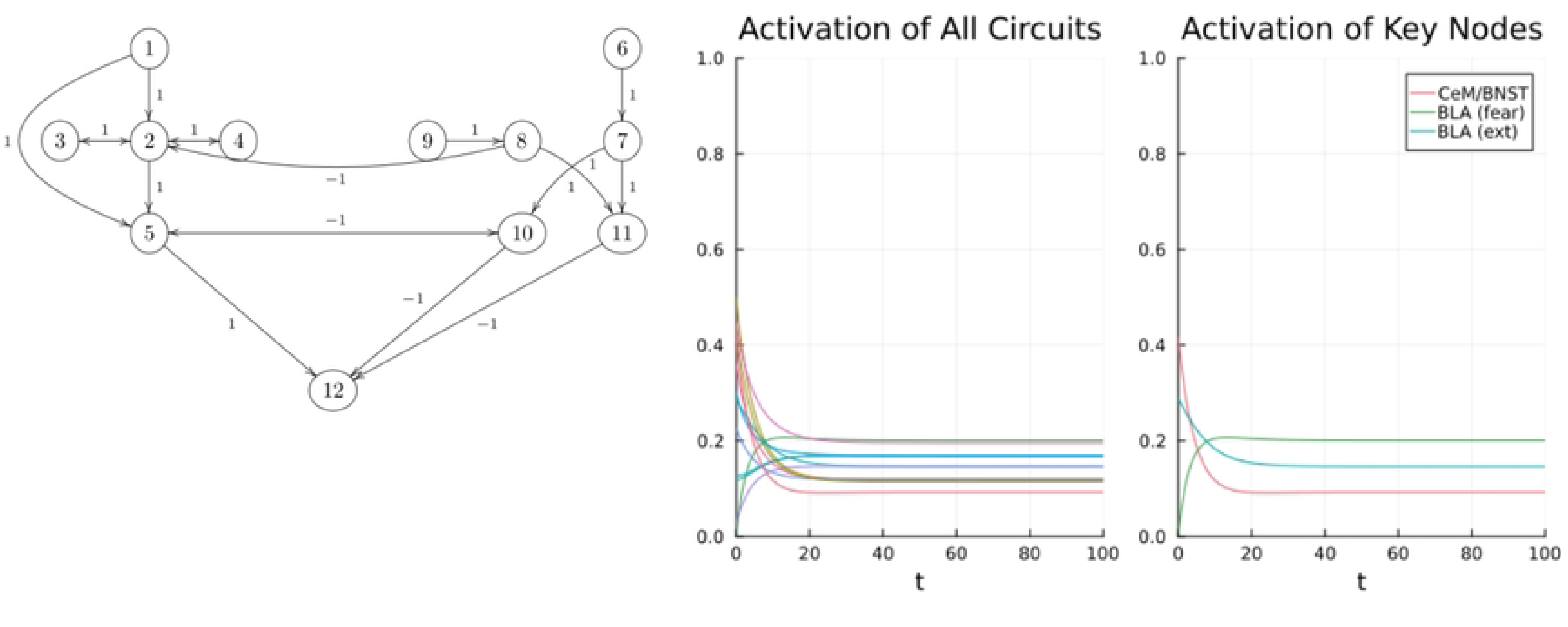
Resting state: circuit, graphs. # 1 =PVTfear, 2 = BLAfear, # 3 = vHPCfear, # 4 = ACC, #5=CeLon. Nodes of extinction are: # 6 =PVText, # 7 =BLAext, #8=vMPFC, # 9 = vHPCext, # 10 - CeLoff, # 11 - ITC, # 12 = CeM/BNST.

### Generalized anxiety disorder, panic disorder, post-traumatic stress disorder

In our model, generalized anxiety disorder (GAD) panic disorder (PD) and post-traumatic stress disorder (PTSD) share the same nodes of the fear and extinction circuits and differ in some aspects of their symptomatology due to degree of stimulation of specific nodes or by the specific activating controlled stimulus (CS) or uncontrolled stimulus (UCS).

For GAD, there is an increase in fear input from PVT fear to BLA fear and a decrease in extinction input from PVText to BLAext. There is also an associated increase in reciprocal fear connections between BLAfear and vHPCfear, and greater CeLon excitation of CM-BNST [90–99]. Stimulation of CM-BNST leads to higher levels of physical arousal through activation of the sympathetic system (increased heart rate, dyspnea, sweating, muscular tension) [92,97]. Stimulation of vACCfear) and vHPC (fear) cause higher psychological symptoms of anxiety (negativity, pessimism, worry, apprehension) [98,99]). Subjects with GAD may not identify a specific CS or UCS. They may have a constant state of apprehension or vigilance toward non-specific threats [98–99]. Levels of symptomatology are governed by moderate levels of activity of BLAfear, vACCfear and vHPCfear. In GAD there is a 25% to 50% increase in nodal activity of the fear-anxiety circuit with a corresponding 25% −75% reduction in activity of extinction nodes, leading to moderate activity of the key anxiety nodes: BLAfear, vHPCfear, ACC and BNST. Human fMRI studies support increased ACC-amygdala activity, increased activity in amygdala, insula, thalamus and posterior cingulate, as well as decreased function of the vMPC [90–96]. Activation of BNST, which causes an increase in activity of the sympathetic nervous system, is present in all anxiety disorders [97–99].

Circuit weightings for GAD are: W[1,2] = 1.25, W[1,5] = 1.5, W[2,3] = 1.5, W[2,4] = 1.25, W[2,5] = 1.25, W[3,2] = 1.5, W[4,2] = 1.25, W[5,10] = −1, W[5,12] = 1.5, W[6,7] = .75, W[7,10] = .75, W[7,11] = .75, W[8,2] = −.5, W[8,11] = .5, W[9,8] = .5, W[10,5] = −.25, W[10,12] = −.25, W[11,12] = −.25

**Fig 2.**
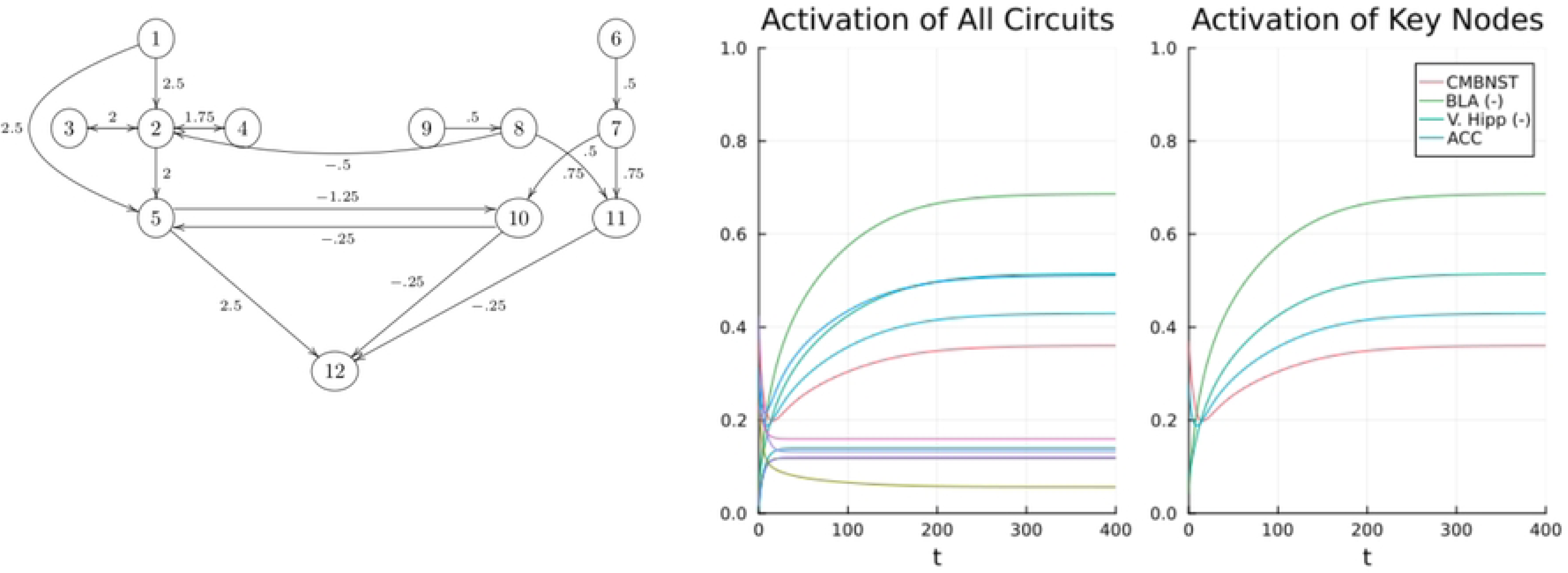
Generalized anxiety disorder, circuits and graphs of nodal activity. # 1 =PVTfear, # 2 = BLAfear, # 3 = vHPCfear, # 4 = ACC, #5=CeLon. Nodes of extinction are: # 6 =PVText, # 7 =BLAext, #8=vMPFC, # 9 = vHPCext, # 10 - CeLoff, # 11 - ITC, # 12 = CeM/BNST.

The hallmark of panic disorder is brief episodes of the highest levels of sympathetic activity. Human neural network data is, of necessity, scant but suggests similarities with non-human studies of severe anxiety states [100–101]. We assume that the neurophysiology of panic attacks in humans resembles that of nonhuman subjects during foot-shock with high activation of the amygdala, hypothalamus, hippocampus, medial prefrontal cortex and PAG [95–99]. There is a brief episode of intense activity of the sympathetic nervous system-high activation of CM-BNST and input to PAG, LC and hypothalamus [98,99].

Panic disorder is usually initiated by a spontaneous panic attack which functions as an unconditioned stimulus (UCS) sensitizing the subject to the environment in which the panic attack occurs which becomes the conditioned stimulus (UCS). Further exposure to the UCS triggers further panic attacks.this process in humans is similar to sensitization and re-exposure in tne non-human model. In our model the CS is assigned a high negative valence by PVTfear and communicated to BLAfear which coordinates its activities with ACC fear and vHPCfear. Inputs from and between extinction nodes are assigned a lower weighting. During a panic attack high levels of psychological anxiety symptoms as well as intense sympathetic activity is triggered by increased activity of fear-anxiety circuit nodes set at 25% −250% greater than baseline and extinction circuit nodal activity is decreased by 25%-50%.

Circuit weightings for panic disorder are: W[1,2]= 2.5, W[1,5] = 2.5, W[2,3] = 1.5, W[2,4] = 1.5, W[2,5] = 2, W[3,2] = 1.5, W[4,2] = 1.25, W[5,10] = −1.25, W[5,12] = 2.5, W[6,7] = .5, W[7,10] = .5, W[7,11] = .5, W[8,2] = −.5, W[8,11] =.75, W[9,8] = .5, W[10,5] = −.5, W[10,12] = −.5, W[11,12] = −.5

**Fig. 3.**
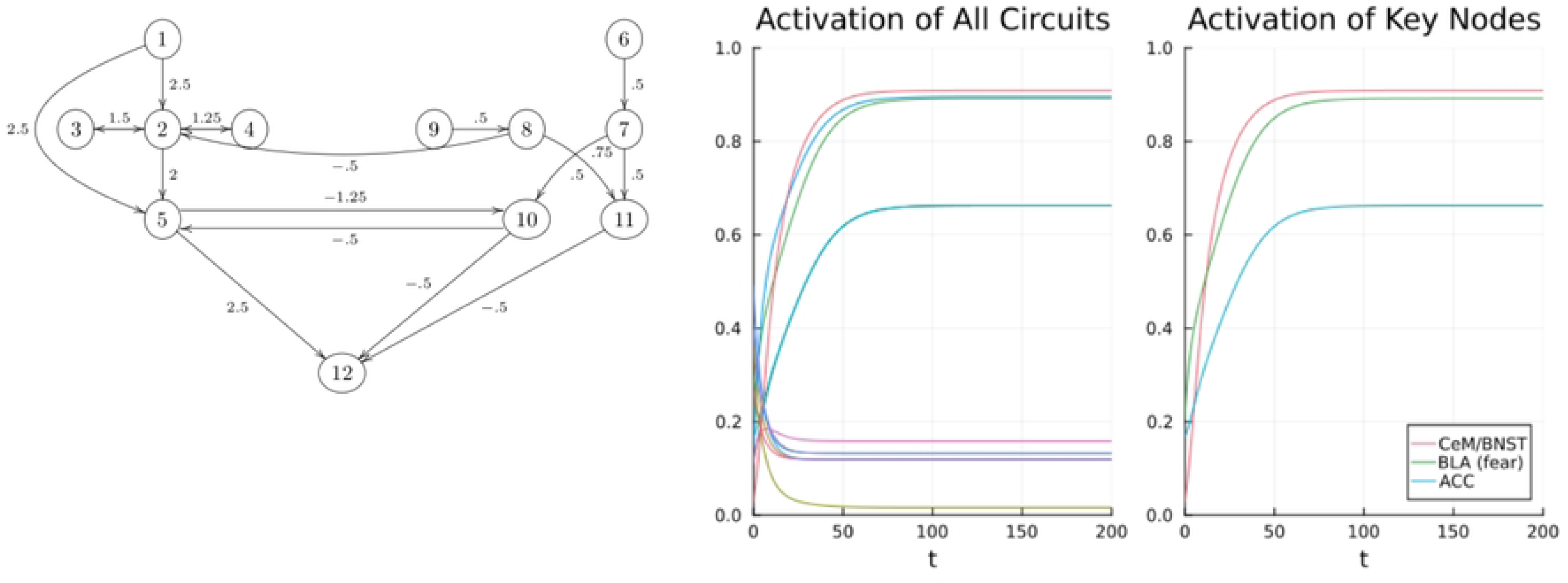
Panic disorder, circuits and graphs of nodal activity during panic attack. # 1 =PVTfear, # 2 = BLAfear, # 3 = vHPCfear, # 4 = ACC, #5=CeLon. Nodes of extinction are: # 6 =PVText, # 7 =BLAext, #8=vMPFC, # 9 = vHPCext, # 10 - CeLoff, # 11 - ITC, # 12 = CeM/BNST.

Subjects with PTSD suffer from moderate to high baseline anxiety characterized by a constant state of vigilance and arousal with episodes of severe sympathetic discharge provoked by a CS which evokes a UCS of severe threat or trauma. Like panic disorder, PTSD is also characterized by episodes of severe sympathetic discharge. Unlike panic disorder, the UCS is an actual severe threat. The graphs are meant to demonstrate the highest levels of PTSD arousal when provoked by CS. The weightings of PVTfear, BLAfear, ACC, and vHPCfear are set higher than in panic disorder to reflect higher negative valence assigned to extreme threat. In human studies hyperactivation of ACC, amygdala, hippocampus, and reduced activity of vMPC are most consistently reported [103–110].

Circuit weightings for PTSD are: W[1,2]= 2.5, W[1,5] = 2.5, W[2,3] = 2, W[2,4] = 1.75, W[2,5] = 2, W[3,2] = 2, W[4,2] = 1.75, W[5,10] = −1.25, W[5,12] = 2.5, W[6,7] = .5, W[7,10] = .5, W[7,11] = .5, W[8,2] = −.5, W[8,11] =.75, W[9,8] = .5, W[10,5] = −.5, W[10,12] = −.5, W[11,12] = −.5.

**Fig. 4.**
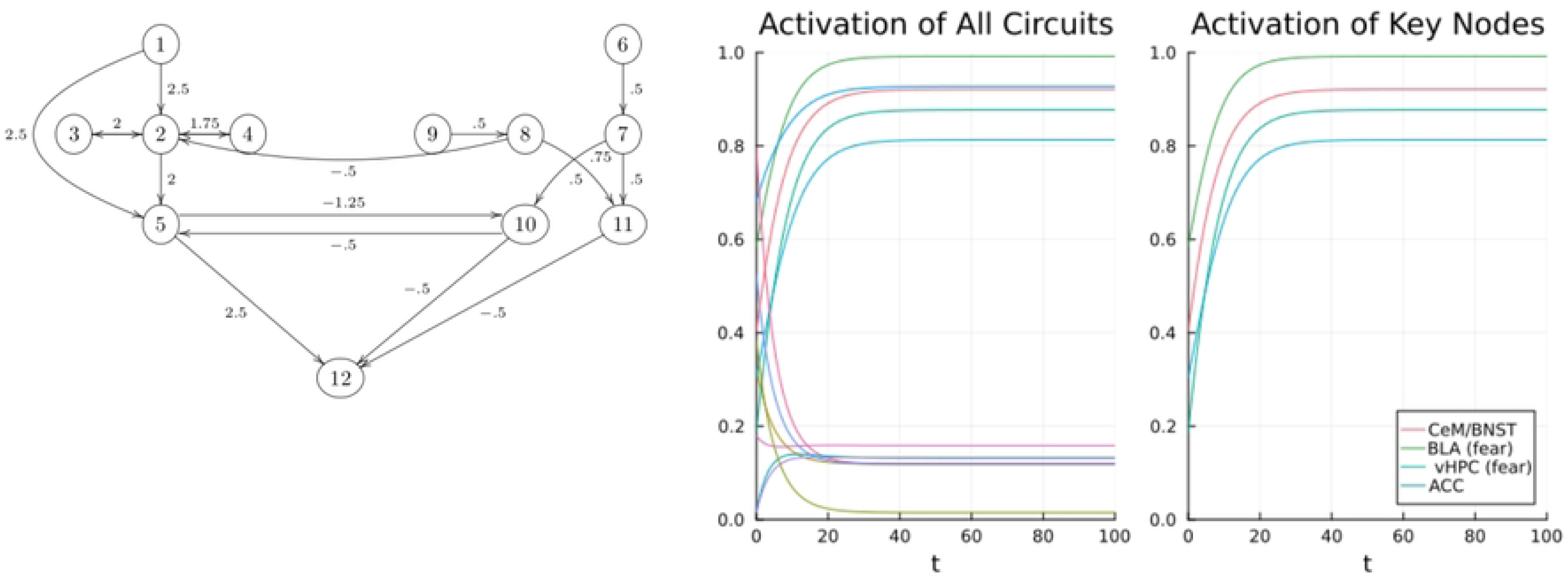
Post-traumatic stress disorder: circuit and graphs of nodal activity during a state of high arousal. # 1 =PVTfear, 2 = BLAfear, # 3 = vHPCfear, # 4 = ACC, #5=CeLon. Nodes of extinction are: # 6 =PVText, # 7 =BLAext, #8=vMPFC, # 9 = vHPCext, # 10 - CeLoff, # 11 - ITC, # 12 = CeM/BNST.

### Social anxiety and obsessive-compulsive disorder

The circuits for social anxiety (SA) and obsessive-compulsive disorder (OCD) include nodes added to the core nodes for GAD, PD and PTSD. For SA, mPFC receives afferents from BLAfear and ACC [111] which produces fear of social encounters and the psychological symptoms of embarrassment and fear of being harshly judged, apprehension, and ruminating over assumed mistakes. The high negative valence assigned to social interactions is communicated directly to the mPFC by BLAfear and ACC, and the vHPCfear stores fear memories of prior humiliations which it communicates indirectly to the mPFC through BLAfear. For SA we must rely more heavily on human data derived from fMRI [112–115]. The circuit weightings for SA includes the mPFC assigned #13.

W[1,2]=1.5, W[1,5] = 1.25, W[2,3] = 1.25, W[2,4] = 1.75, W[2,5] = 1.25, W[2,13] = 1.5, W[3,2] = 1.25, W[4,2] = 1.75, W[4,13] = 1.5, W[5,10] = −1.25, W[5,12] = 1.5, W[6,7] = .5, W[7,10] = .75, W[7,11] = .75, W[8,2] = −.75, W[9,8] = .5, W[10,5] = −.75, W[10,12] = −.75, W[11,12] = −.75 .

**Fig. 5.**
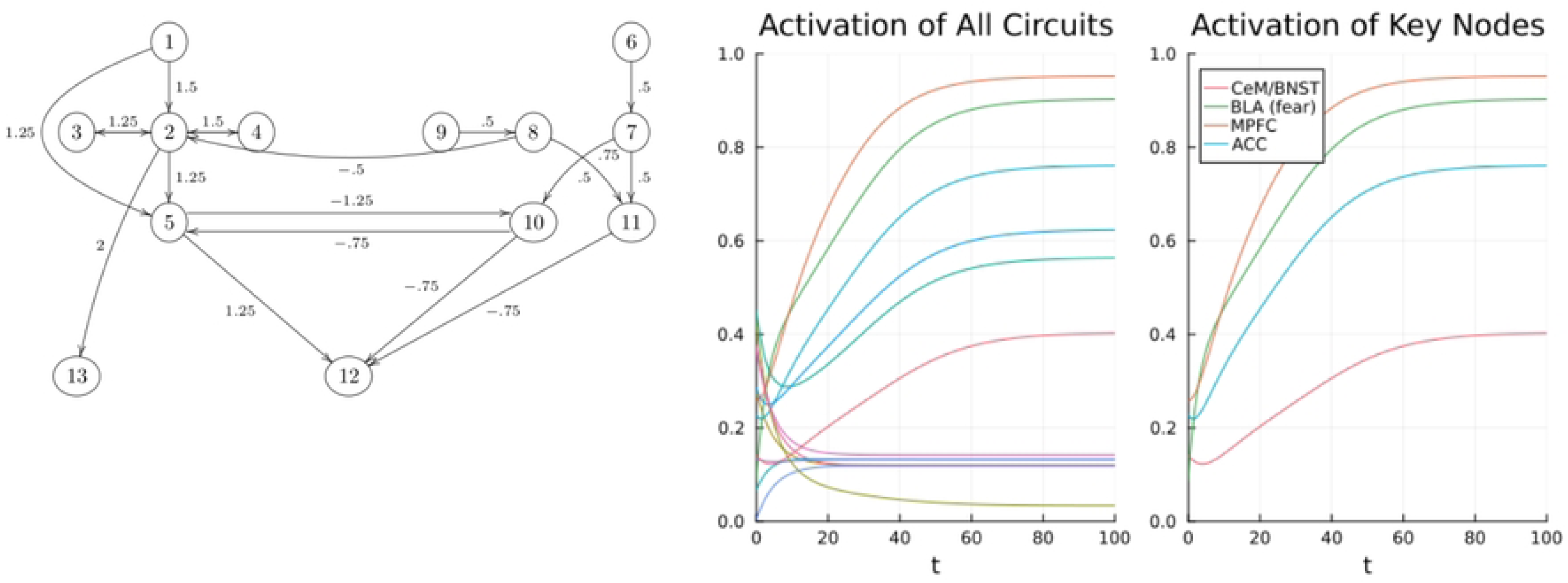
Social anxiety disorder, circuit and graphs of nodal activity. # 1 =PVTfear, 2 = BLAfear, 3 = vHPCfear, # 4 = ACC, #5=CeLon. Nodes of extinction are: # 6 =PVText, # 7 =BLAext, #8=vMPFC, # 9 = vHPCext, # 10 - CeLoff, # 11 - ITC, # 12 = CeM/BNST, #13=MPC.

OCD has the most complex fear circuits underlying the psychological symptoms of obsessions and the behavioral symptoms of compulsions. For OCD we must rely on data from human fMRI studies along with nonhuman animal studies which most consistently identify hyperactivity of amygdala, orbitfrontal cortex, ACC and ventral striatum as well as hypoactivity of vmPFC [115–122]. In our model, the characteristic symptoms of OCD are produced by strong negative valence signals from BLAfear to OFC and Str as well as strong fear stimulation of ACC by both BLAfear and OFC [115–132]. The activity of OFC produces our sense of rightness in general and our social constructs of morality or transgression. A persistent negative valence signal to the OFC causes obsessive thoughts [116–118, 122–124]. The striatum constructs behavioral repertories: A negative valence signal directed to the striatum leads to elevated levels of repetitive activities which the subject feels are compelled [125–131]. Circuit weightings for OCD include Str assigned #14 and OFC assigned #15.

W[1,2] = 2, W[1,5] = 2, W[2,3] = 2, W[2,4] = 2, W[2,5] = 2, W[2,13] = 1, M[2,14] = 1, W[3,2] = 2, W[4,2] = 2, W[5,10] = −1.5, W[5,12] = 1.75, W[6,7] = .5, W[7,10] = .5, W[7,11] = .5, W[8,7] = .5, W[8,11] =.5, W[9,8] = .5, W[10,5] = −.5, W[10,12] = −.5, W[11,12] = −.5, W[2,15]= 2.0, W[15,4]= 2.0, W[15,14]= 2.0.

**Fig. 6.**
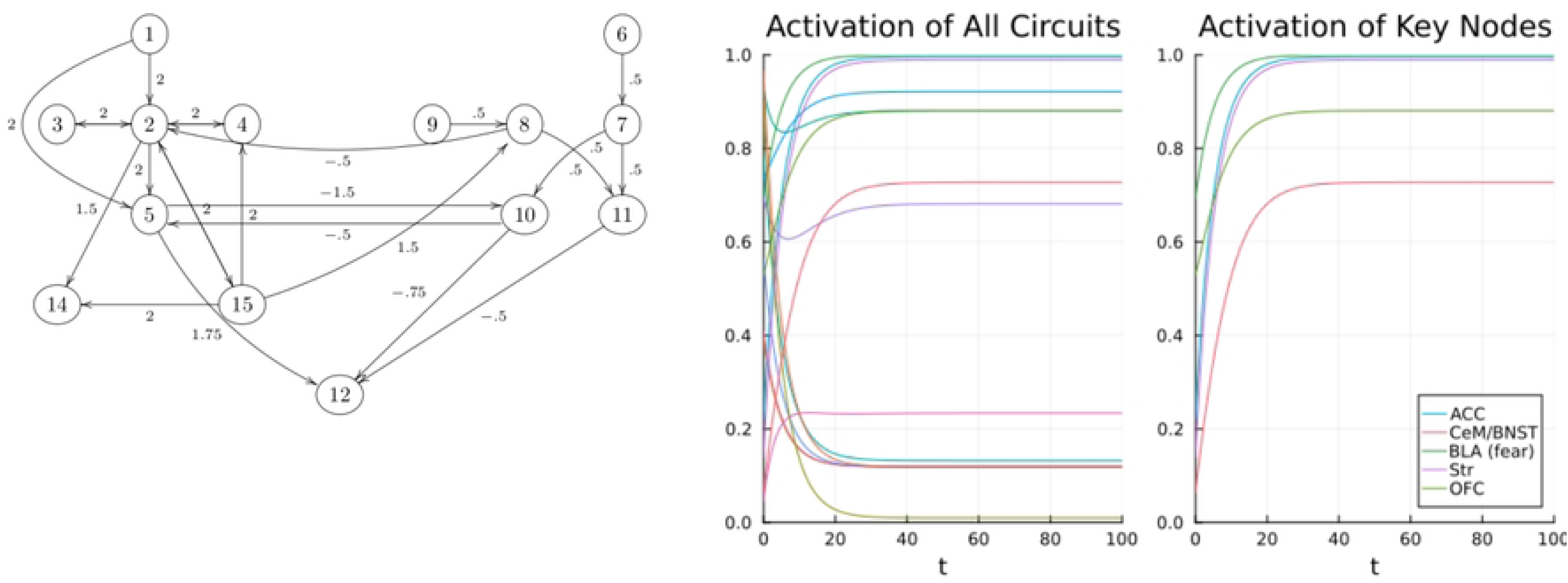
Obsessive compulsive disorder, circuits, graphs of nodal activity. # 1 =PVTfear, 2 = BLAfear, # 3 = vHPCfear, # 4 = ACC, #5=CeLon. Nodes of extinction are: 6 =PVText, # 7 =BLAext, #8=vMPFC, # 9 = vHPCext, # 10 - CeLoff, # 11=ITC, # 12 = CeM/BNST, #13=MPC, #14=Str, #15=OFC.

## Discussion

The main findings of this paper are that the symptom clusters of anxiety disorders, their origin and extinction can be explained by neural network theory. We have presented what we believe is a unique formal model of anxiety disorders arising from the dynamics of coupled fear and extinction circuits.

The preservation of limbic structures allows for the precise neurophysiologic exploration of fear and extinction circuits in non-human animal models for the insights unavailable in human studies. Human fMRI studies in anxious human subjects offer less precise information on the activity of specific nodes of the human fear and extinction networks due to lack of sufficient resolution.

Non-human studies suggest a common substrate for the generation and expression of core anxiety symptoms. and confirm the centrality of the nodes found in anxiety disorders. GAD, SA, PD, and PTSD have highly analogous animal model symptoms provoked by similar stresses as in human subjects [14–22]. We suggest that SSRIs and therapeutic techniques based upon extinction are effective for both humans and nonhuman subjects because the same symptom clusters arise from similar neural pathways of fear and extinction which have been highly conserved across species [132–136].

Our model assumes that anxiety disorders are the result of deficiencies of BDNF function due to genetic or epigenetic alterations in BDNF or its receptors [61–87]. Genetic and epigenetic studies in both human and non-human species confirm low BDNF or decreased BDNF receptors in key nodes of the extinction circuit [69–79]. Our neural circuit model of anxiety pathophysiology proposes that genetic alterations or stress-induced methylation of the BDNF genes cause pathologic function of the extinction circuit [72–79] which leads to anxiety disorders.

Our formal model explains how low BDNF alters the function of key nodes by reducing the neuroplasticity needed to produce extinction responses believed to underly the symptom clusters pathognomonic for anxiety disorders. The decrease in synaptic plasticity is represented by a decrease in connectivity (*W*) between extinction nodes and their efferents. Serotonin receptor inhibitors are effective for anxiety disorders by increasing BDNF function of affected nodes of the extinction circuit [132–136].

We believe that further gains in the understanding of anxiety disorders will arise by employing the analytic rigor and precision of mathematics to the dynamic interactions of the fear-anxiety and extinction circuits and their constituent nodes.

Knowledge of neural circuits has accumulated at the same time as the development of increasingly exact brain imaging techniques combining fMRI with diffusion tensor imaging or MRI. Progress is also being made in the ability to deliver a more discreet targeted electromagnetic field to deep structures of the brain using non-invasive deep transcranial temporal interference stimulation [137–141].

Though speculative and qualitative, we offer a theoretical model for the accumulation of more data which will ultimately lead to a more definitive and quantitative version. We believe that our formal model of anxiety disorders may offer a heuristic insight into the further development of innovative treatments.

## Acknowledgements

None

